# Machine learning enables accurate prediction of breast cancer five-year survival using somatic genomic variants

**DOI:** 10.1101/2022.05.22.492994

**Authors:** Xiaosen Jiang, Laizhi Zhang, Guangshuo Cao, Jia Li, Yong Bai

**Author notes:** Correspondence: Yong Bai.

## Abstract

Breast cancer is one of the most common cancers, accounting for about 30% of female cancers and a mortality rate of 15%. The 5-year survival rate is most commonly used to assess cancer progression and guide clinical practice. We used the CatBoost model to systematically construct a five-year mortality risk prediction model based on two independent data sets (BRCA_METABRIC, BRCA_TCGA). The model input data are the somatic genomic variants (copy number variation, SNP locus, cumulative mutation number of genes) and phenotype data of cancer samples. The optimal model combined all the above characteristics, and the AUC reached 0.70 in an independent external data set. At the same time, we also conducted a biological analysis of the characteristics of the model and found some potential biomarkers (TP53, DNAH11, MAP3K1, PHF20L1, etc.). The results of model risk stratification can be used as a guide for the prognosis of breast cancer.

## Introduction

Breast cancer has overtaken lung cancer to become the most common malignancy and the first leading lethal cancer among women, accounting for approximately 11.7% of all cancer cases diagnosed (2.3 million) in 2020 around the world[1]. In spite of significant medical improvements in early diagnosis and modern therapy [2], breast cancer still poses increasing burden globally and the overall survival outcomes remain unsatisfactory given the mortality of 1 in 6 female cancers (685,000 deaths)[1], reflecting high biological complexity and genetic heterogeneity of the disease[3-5]. Consequently, identification of genetic prognostic indicators plays crucial roles in understanding inter-individual differences in pathogenesis between breast cancer patients, providing better insight into therapeutic decision-making and optimizing personalized precise treatment. Additionally, the prognosis of breast cancer, which has a 5-year survival rate greater than 85%, is better than other cancers[6]. Since the threshold of five years is most commonly used to assess the process of the cancer, we can anticipate the 5-year mortality of breast cancer to guide clinical intervention.

One of the earliest survival prognostic models, Nottingham Prognostic Index(NPI)[7], was constructed based on clinical factors using Cox regression[8]. Since then, more variables had been considered to improve the accuracy of NPI[9]. In 2001, Adjuvant, an internet-based tools was developed and widely applied in prognosis analysis in breast cancer[10]. In 2010, two independent models, OPTIONS[11] based on parametric regression and PREDICT based on Cox proportional hazards, were established. But both Adjuvant and PREDICT showed low accuracy of estimating mortality in different sub-groups, especially for young breast cancer patients[12]. Additionally, these statistically constructed models based on clinical data are limited by the prolonged process of data collecting and poor timeliness. These drawbacks entail the need for prediction using data collected right after diagnosis. With the advent of microarray-based gene expression profiling, some gene-related studies demonstrate the impact of genetic factors on prognostic and survival prediction of breast cancer and lots of predictive signatures have been found[13]: such as MammaPrint[14], Oncotype DX[15], Endopredict[16]. Although these models are applied to sub-groups of breast cancer patients, their signatures cannot sensibly interpret their relationships with breast cancer outcomes, which are called black-box models[17]. Thus models with high accuracy and high interpretability need to be further developed.

Machine learning(ML) is a feasible method, since ML can extract features from large genetic datasets and perform risk scoring and classification[18]. Genetic signature copy number alteration(CNA) has a strong correlation with the prognosis and mortality of cancer[19]. However, because the genomic CNA dataset is large and relatively sparse, traditional models based on single or several CNA signatures are not explicable. Furthermore, Somatic mutations can also be used to construct predictive models for risk scoring and survival prediction. Nguyen et al. selected multi-features with Random Forest(RF), largely improving the accuracy of the predictive model[20]. Support Vector Machine(SVM), Artificial Neural Network(ANN) and semi-supervised learning methods were employed to construct predictive models for assessing the survivability of breast cancer patients[21]. Compared to the integrated model, the models based on somatic mutations alone have lower accuracies, and integrated models are limited by small samples and may have overfitting problems[22, 23]. We found that most of the reported 5-year survival prediction models for breast cancer have considered data preprocessing, feature selection, class imbalance processing, and model validation.[24]. We only find two studies that were further verified externally[25, 26]. Both of the studies used the Molecular Taxonomy of Breast Cancer International Consortium(BRCA_METABRIC) dataset for training and internal validation, and The Cancer Genome Atlas(BRCA_TCGA) dataset was used for further validation. However, after scrutinizing these two studies, we found that their independent dataset, which should be used for external test, was fed into their models for training and internal testing again. External validation is necessary because it can reflect the generalization ability of the predictive model. So far as we know, there is no breast-survival-predictor that undertakes external test using independent cancer datasets.

In our study, we developed a CatBoost-based machine learning model that integrates multi-dimensional data including single-nucleotide variants(SNV), cumulative number of gene mutations(CNGM), copy number alteration(CNA) and phenotype data. BRCA_METABRIC dataset was employed for training and internal validation and BRCA_TCGA dataset for external testing. The final result of the model has good generalization ability, and the AUC in the external test set is 0.70. In addition, the feature interpretation of the model found that the model has a high learning ability, and some features that have been reported to be highly related to cancer were found in the model. The code required by the model can be viewed in github: https://github.com/jxs1996/Breast_cancer-5-year-survival-prediction

## Materials and methods

### Data preparation

We obtained two breast cancer data sets from the public database cBioPortal(BRCA_TCGA(n=1108 samples)[27], https://www.cbioportal.org/study/summary?id=brca_tcga; BRCA_METABRIC(n=2509 samples) [28-30], https://www.cbioportal.org/study/summary?id=brca_metabric). Overall, the median age was 62□years; the 5-year survival rate was 75%. Both data sets contain clinical data, somatic mutations, CNA and gene expression data. In the preliminary data processing(Figure1.A), The SNV, CNGM, and CNA features in the two data sets were separately counted and constructed into the input data required by the model. Predictive features were expressed as follows: (1)SNV: if there is a mutation, it is marked as “1”, if it is not marked as “0”; CNGM: each additional SNV, the value increases by 1, and each additional insertion or deletion, the value increases by 10 (This is because we assume that insertions and deletions have a greater accumulation of mutations and a greater impact on genes than SNV); CNA: “-2” for homozygous deletion, “-1” for hemizygous deletion, “0” for neutral / no change, “1” for gain, “2” for high level amplification. The missing values of all features are filled with “0”. Gene expression data is not included in the feature representation. There are two main reasons. 1) The data obtained from the two datasets are standardized with Z-score, which means they are not in the same spatial dimension. Training results in one data cannot be applied in another dataset; 2) If genomic data alone yields better prediction results.This can significantly reduce the cost of the application. Nevertheless, gene expression data will be used in subsequent analysis to observe the performance of the model features at the transcriptome level. In order to ensure the consistency of features, the intersection of SNV, CNGM and CNA is obtained in two independent datasets. Next, we labeled samples as survival(OS_Months > 60) or death(OS_MONTHS < 60 and OS_Status = deceased). Some sample data were discarded(OS_MONTHS < 60 and OS_STATUS = LIVING).

**Figure 1.**
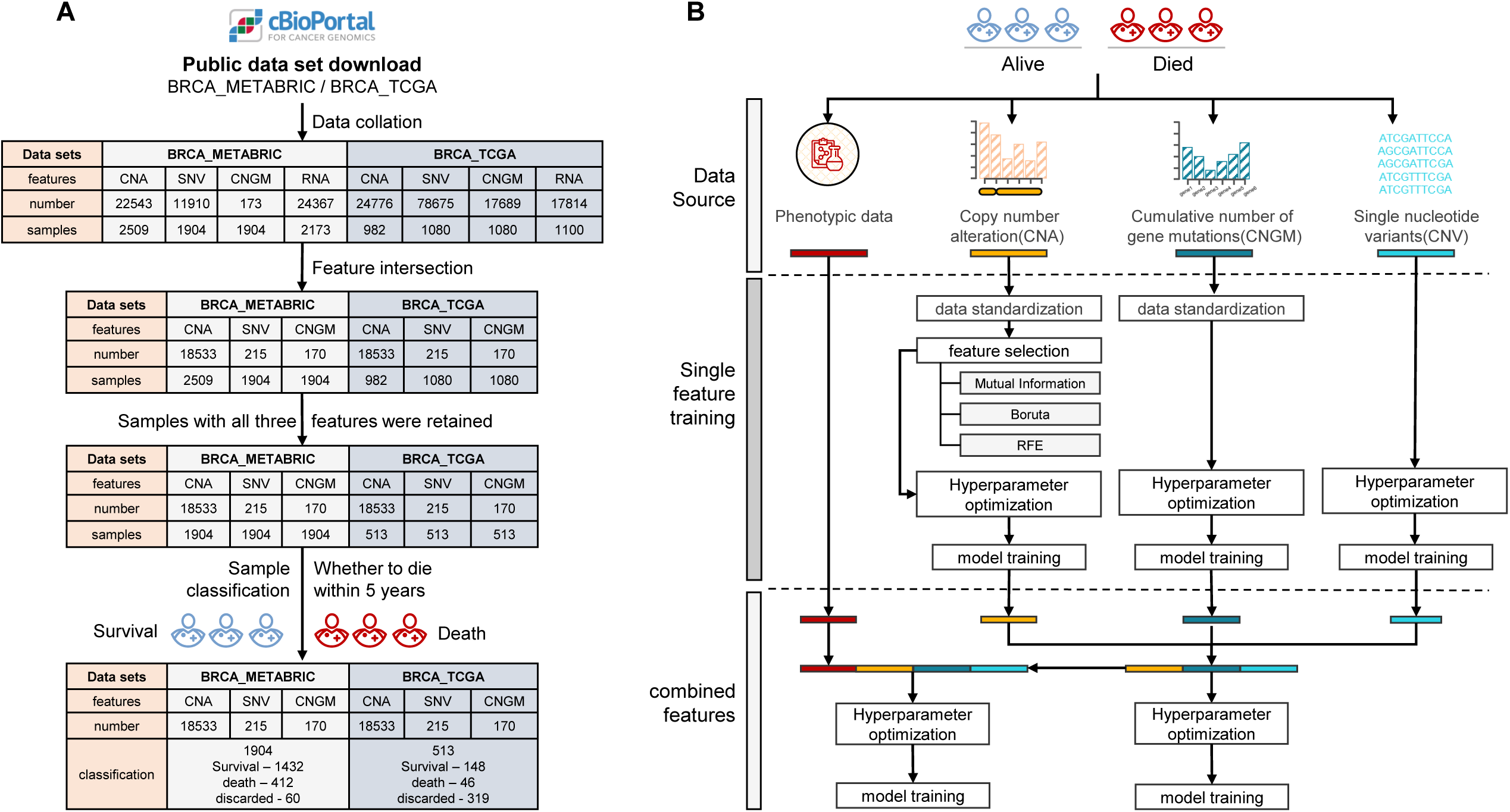
Data quality control and five-year survival prediction model building process. **A)** BRCA_TCGA and BRCA_METABRIC data acquisition and quality control process; **B)** In the process of building a polygenic risk assessment model, different processing methods are adopted for different dimension characteristics.

We retained a total of CNA - 18533, SNV - 215, CNGM - 170 in both datasets. BRCA_METABRIC dataset retained 1904 individuals(survival - 1432, death - 412, discarded - 60), BRCA_TCGA dataset retains 513 individuals(survival-148, death-46, discard-319). In addition, we screened the clinical data shared by the two datasets(because we hope to establish an early risk prediction model, the data of intervention treatment will not be considered), and finally obtained age, gender, number of positive limph nodes and menopausal state. Most patients are female (there are only three males in the BRCA_TCGA dataset), so the sample is no longer grouped by gender. The statistical results of other phenotypes are shown in Supplementary Figure S1. The average age of all breast cancer patients is 60.65 years, and the average lymph node is 2.02. 443 are not yet menopausal, and 1548 are in menopause. If there is no measurement data in the phenotype, it will be represented by -9 and will not participate in the mean calculation.

### Machine learning analysis process development

As shown in Figure 1.B, Catboost, a high-performance open source library for gradient boosting on decision trees, was developed to predict the five-year mortality risk of patients. The analysis process is systematically constructed using the machine learning framework scikit-learn(https://scikit-learn.org/stable/, version=0.24.2). The BRCA_METABRIC data(1844 samples: survival-1432, death-412) set was split into training set(80%) and testing set(20%) by random stratified sampling. The independent external data set BRCA_TCGA (194 samples: survival-148, death-46) will be used for model evaluation. For the three features(SNV, CNA, CNGM), separate models are established to evaluate their effects on prediction. After that, we extract the model features constructed by SNV, CNA, and CNGM and merge them to construct a new multi-dimensional feature set for training and evaluation. Finally, phenotypic characteristics (age, number of positive lymph nodes, menopausal status) are also integrated to further improve the accuracy of the model.

### Single-dimensional feature model construction

Separate models were constructed for CNV, SNV, and CNGM characteristics to explore their impact on the five-year mortality risk prediction. As shown in Figure 1.B, For CNA (18533 features), the training set is first standardized (StandardScaler method), the average value and standard deviation are retained and then applied to the corresponding features in the test phase, and feature selection is performed on the processed data (described in Feature selection part). Next, use the CatBoost model to select the hyperparameters of the model (described in Hyperparameter selection part). After fixing the hyperparameters, perform model training. Five-fold cross-validation is used to evaluate the stability of the model, and finally tested on the test set and independent external data set. The processing of SNV and CNGM is similar to CNA, but due to the small number of SNV features (215) and onehot-encoded, data standardization and feature selection are not performed; CNGM uses log first and then logMinMaxScaler method during standardization. Since there are only 170 features, feature selection is also omitted.

### Multi-dimensional feature model construction

Combine the feature selection results of the single-dimensional feature model and perform hyperparameter selection (described in Hyperparameter selection part). After the hyperparameter results are fixed, perform training and evaluation. In addition, we combined the phenotypic data (age, gender, number of positive lymph nodes, and menopausal status) with the feature selection results of the single-dimensional feature model to observe whether the phenotypic data can improve the performance of the model.

### Feature selection

For CNA(18533 features), irrelevant features may decrease the performance of the model. We propose a hybrid feature selection method to subtract features. In this method, mutual information (MI) technology[31], recursive feature elimination (RFE) algorithm[32] and Boruta algorithm[33] are used to obtain the relevant subset of the raw features. MI calculates feature weights based on the relationship between features of mutual information; RFE selects features by recursively considering smaller and smaller feature sets, and this method can obtain all feature rankings. The Boruta algorithm is a packaging method that selects a subset of features based on a random forest machine learning algorithm, which can be used to measure the importance of features. Respectively use the above methods to obtain the feature ranking and retain the top 3% features (extract the most effective features and maintain a balance with the SNV and CNGM feature numbers). The features selected by any two methods will be retained eventually.

### Hyperparameter selection

Due to the imbalance between death and survival samples (∼1:3), when 80% of the training set is used for hyperparameter training, a small number of samples are randomly sampled to make the ratio of positive and negative samples reach 1:1. We implemented a basic grid search algorithm with 5-fold cross-validation to optimize the Catboost model parameters while maximizing the weighted F1 score.

### Model comparison

After the model training is completed, we will use the five-fold cross-validation data set, test set and independent external data set to evaluate the model. The specific evaluation indicators are as follows:

*TP:* True Positive. In the samples judged to be positive, the number of correct judgments.

*FP:* False Positive. In the samples judged as positive, the number of judgment errors.

*TN:* True Negative. Among the samples judged as negative, the correct number is judged.

*FN:* False Negative. In the samples judged as negative, the number of judgment errors.

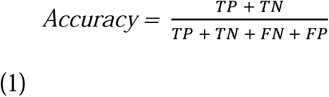

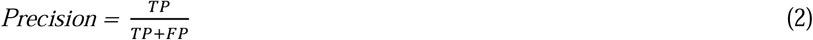

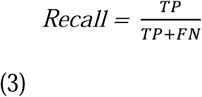

*AUC:* The area under the receiver operating characteristic(ROC) curve, is currently considered to be the standard method to assess the accuracy of predictive distribution models[34]. with AUC = 1 represents perfect performance and 0.5 means random guess.

*F1-score:* The harmonic mean of the precision(2) and recall(3). The highest possible value of an F-score is 1.0, indicating perfect precision and recall, and the lowest possible value is 0, if either the precision or the recall is zero.

### Feature analysis

We will select the model with the best comprehensive score and use the SHAP(Shapley Additive exPlanations) tool to analyze the characteristics of the final model[35]. SHAP is a unified method to explain machine learning predictions based on the optimal Shapley value of game theory. SHAP computed the contribution of each feature to the prediction, which was quantified using Shapley values from coalitional game theory. The Shapley value was represented as an additive feature attribution method, providing the average of the marginal contributions across all permutations of features and distribution of model prediction among features. As an alternative to permutation feature importance, SHAP feature importance was based on magnitude of feature attributions. The absolute Shapley values per feature across the data was further averaged as the global importance was needed. We ranked the features importance in descending order and picked the top 30 most important features. The SHAP value can be plotted for each sample corresponding to the first 30 features. We used the Python library to implement the SHAP algorithm (https://github.com/slundberg/shap).

For features, the enrichment analysis in CLINVAR, KEGG, GO, and Reactome will also be performed using ClueGO[36]. In addition, the genes corresponding to the optimal model features were extracted, and the Kruskal-Wallis test was used in the BRCA_METABRIC gene expression data set (Bonferroni correction of the results, adjusted P value <=0.05) to calculate the difference between survival and death groups. For genes with significant differences, use the limma tool[37] to calculate the expression fold difference.

### Risk stratification analysis

The original output result of the model is a probability value (between 0 and 1). Based on the optimal model result, We will divide all samples into high, medium and low risk groups (BRCA_METABRIC, BRCA_TCGA), and draw Kaplan-Meier (K-M) curve.

## Result

### Comparison of performance of different machine learning models

The optimization process of the five models (CNA, SNV, CNGM, SNV+CNGM+CNA(combined variants), combined variants+phenotype)was shown in Supplementary Figure S2. The number of features and AUC values corresponding to the optimal model were: SNV(AUC:0.56; features: 93), CNGM(AUC:0.63; features: 4), CNA(AUC:0.64; features: 75), combined variants (AUC:0.72; features: 353), combined variants + phenotype (AUC:0.81; features: 172).

We have drawn the Precision-Recall and ROC curves for the above optimal models using the 5-fold cross-validation method in the BRCA_METABRIC, internal test set and external test set (Figure 2). Taking the test result of the external data set BRCA_TCGA as the final evaluation index, the indexes of each model are as follows: SNV(AUC:0.53, APS:0.25); CNGM(AUC:0.54, APS:0.26); CNA(AUC:0.62, APS:0.42); combined variants (AUC:0.61, APS:0.35); combined variants+phenotype(AUC:0.70, APS:0.43); More model evaluation indicators can be viewed in Table 1. The best comprehensive score was the combined variants+phenotype model, which performed best in both the internal test set (AUC:0.81, APS:0.55) and the independent external data set (AUC=0.70, APS:0.43)(Table 1).

**Figure 2.**
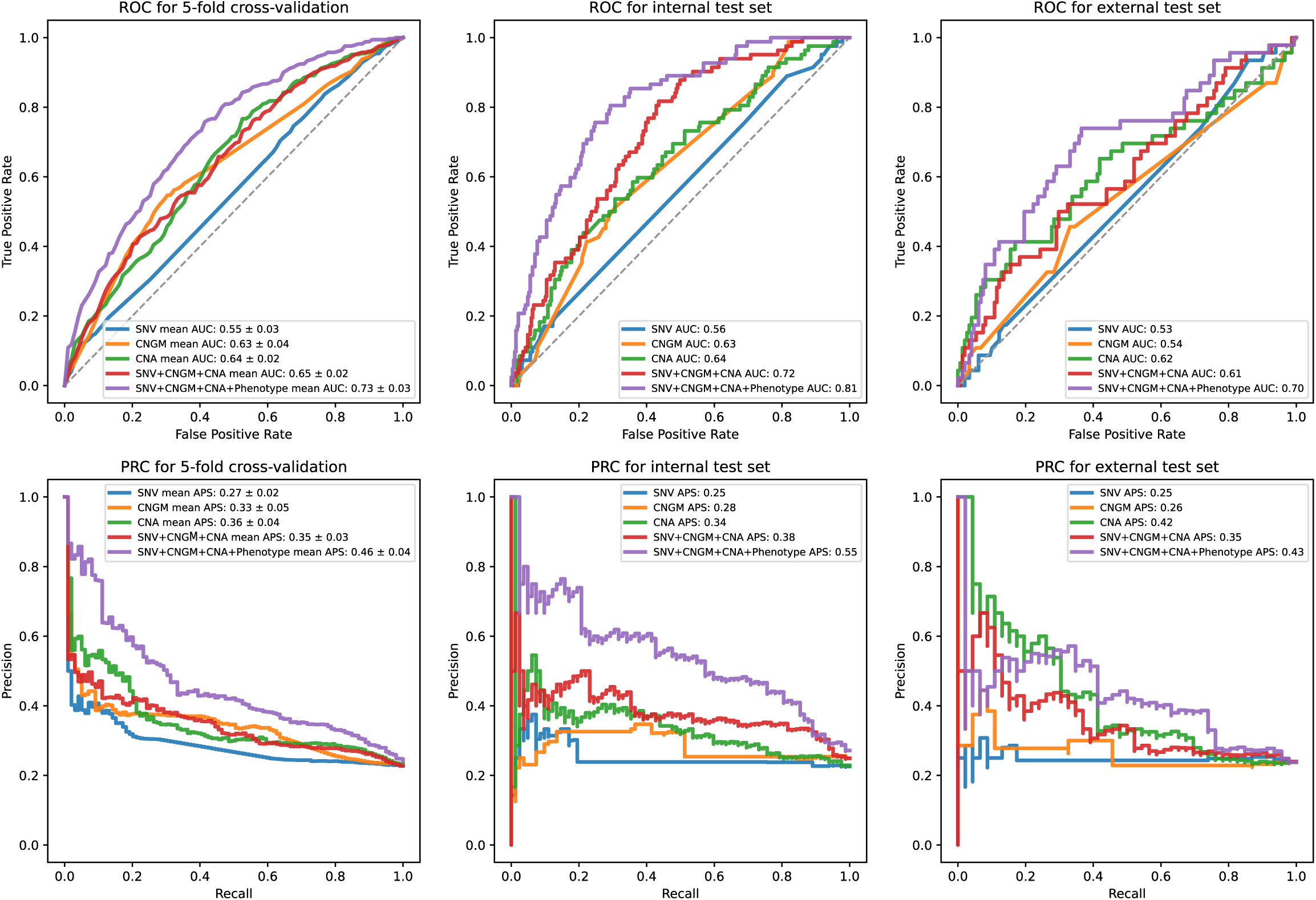
Precision-Recall and ROC curves of optimal models constructed with features of different dimensions. The first row is the ROC curve and the second row is the Precision-Recall curve. From left to right are the results in the cross-validation set(mean), training set and test set, respectively.

**Table 1.** The model predicts the performance indicators of breast cancer deaths within five years in the internal and external test data sets.

| Model | Internal test data |  |  |  |  |  | External test data |  |  |  |  |  |
| --- | --- | --- | --- | --- | --- | --- | --- | --- | --- | --- | --- | --- |
|  | AUC | F1 | Accuracy | precision | recall | APS | AUC | F1 | Accuracy | precision | recall | APS |
| SNV | 0.56 | 0.21 | 0.72 | 0.29 | 0.17 | 0.25 | 0.53 | 0.22 | 0.70 | 0.29 | 0.17 | 0.25 |
| CNGM | 0.63 | 0.37 | 0.66 | 0.32 | 0.45 | 0.28 | 0.54 | 0.36 | 0.62 | 0.30 | 0.46 | 0.26 |
| CNA | 0.64 | 0.34 | 0.74 | 0.39 | 0.30 | 0.34 | 0.62 | 0.38 | 0.77 | 0.52 | 0.30 | 0.42 |
| SNV+CNGM+CNA | 0.72 | 0.32 | 0.76 | 0.42 | 0.26 | 0.38 | 0.61 | 0.25 | 0.73 | 0.36 | 0.20 | 0.35 |
| SNV+CNGM+CNA<br>+Phenotype | 0.81 | 0.52 | 0.80 | 0.55 | 0.49 | 0.55 | 0.70 | 0.46 | 0.77 | 0.51 | 0.41 | 0.43 |

### Optimal model feature ranking

The combined variants + phenotype model comprised a total of 172 features, including 121 CNA, 45 CNGM, 4 SNV, and 2 phenotypes (age, number of positive limph nodes). We used shap to analyze the importance of the predictive characteristics of the model. As shown in Figure 3, among the 172 features of the model, we extracted the top 30 most important features. The phenotypic characteristics age and number of positive lymph nodes ranked first and second, and showed positive correlation with death within five years. The remaining 28 features included 18 CNA (ZNF720, TBC1D13, SCAF4, CDRT15, TMED6, OR4M2, C17orf102, TAS2R10, PHF20L1, RNF187, STIM2, CCDC136, TTI2, MTBP, FAM24B, TMEM26, OR4F15, PDCL2), 9 CNGM (TP53, DNAH11, DNAH2, PIK3CA, MAP3K1, GATA3, CDH1, PDE4DIP, 80273), 1 SNV(chr3:178936091:G:A). Some characteristics also showed positive correlation with mortality within five years, such as CNGM-TP53, CNGM-DNAH2, CNGM-PIK3CA, CNA-SCAF4, CNGM-CDH1, etc. There were also some opposite manifestations, such as CNA-ZNF720, CNGM-DNAH11, CNA-TMED6, CNGM-MAP3K1, SNP-3-178936091-G-A, etc.

**Figure 3.**
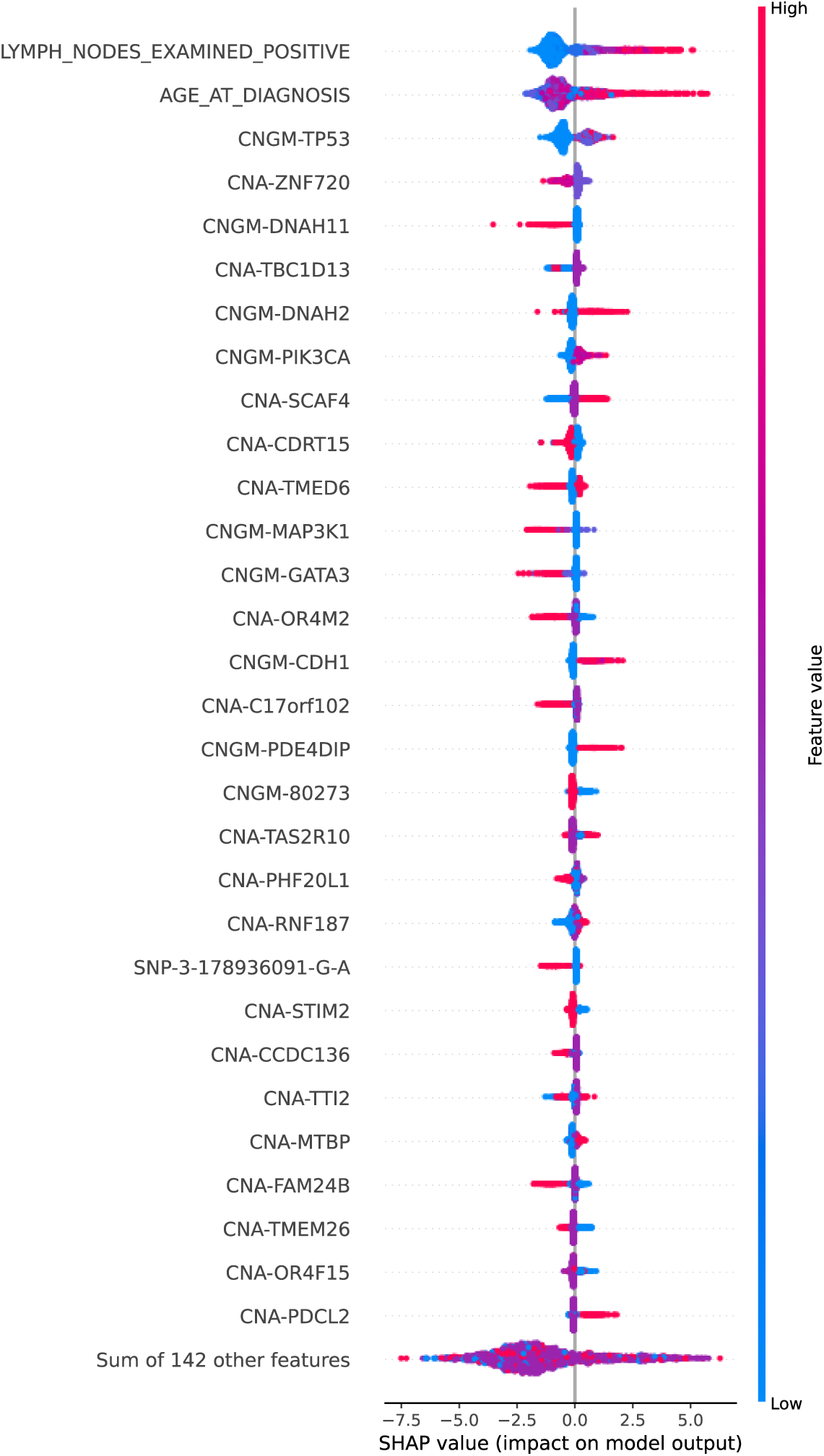
Optimal model feature weight analysis. The scatter points represent the SHAP value of each feature for each sample. Features are sorted according to the sum of the magnitudes of the SHAP values of all samples. The first 30 features are shown, and the colors represent the feature values (red high, blue low). For example, as age (“AGE_AT_DIAGNOSIS”) increases, the risk of death within five years of the sample will increase.

### Enrichment analysis of optimal model features

We used ClueGO to perform enrichment analysis on the genes corresponding to 172 features. The selected data sets included CLINVAR, KEGG, GO, and Reactome pathways. The enrichment results were corrected by bonferoni multiple test. After correction, the pathways with adjusted P value less than 0.05 were selected(Figure 4.A). A total of 33 records were obtained. In CLINVAR and KEGG, the features were enriched in pan-cancer or breast cancer-related pathways (C0006142, C1458155, KEGG:05212, KEGG:05222, KEGG:05224). In GO biological process pathways, these genes were over-represented in some pathways related to cell cycle and cell proliferation (GO:0048103, GO:1904030, GO:0000079, GO:0061982, etc.). Two REACTOME pathways reached statistical significance, both of which are related to the NOTCH signaling pathway (R-HSA: 350054, R-HSA: 1980143)

**Figure 4.**
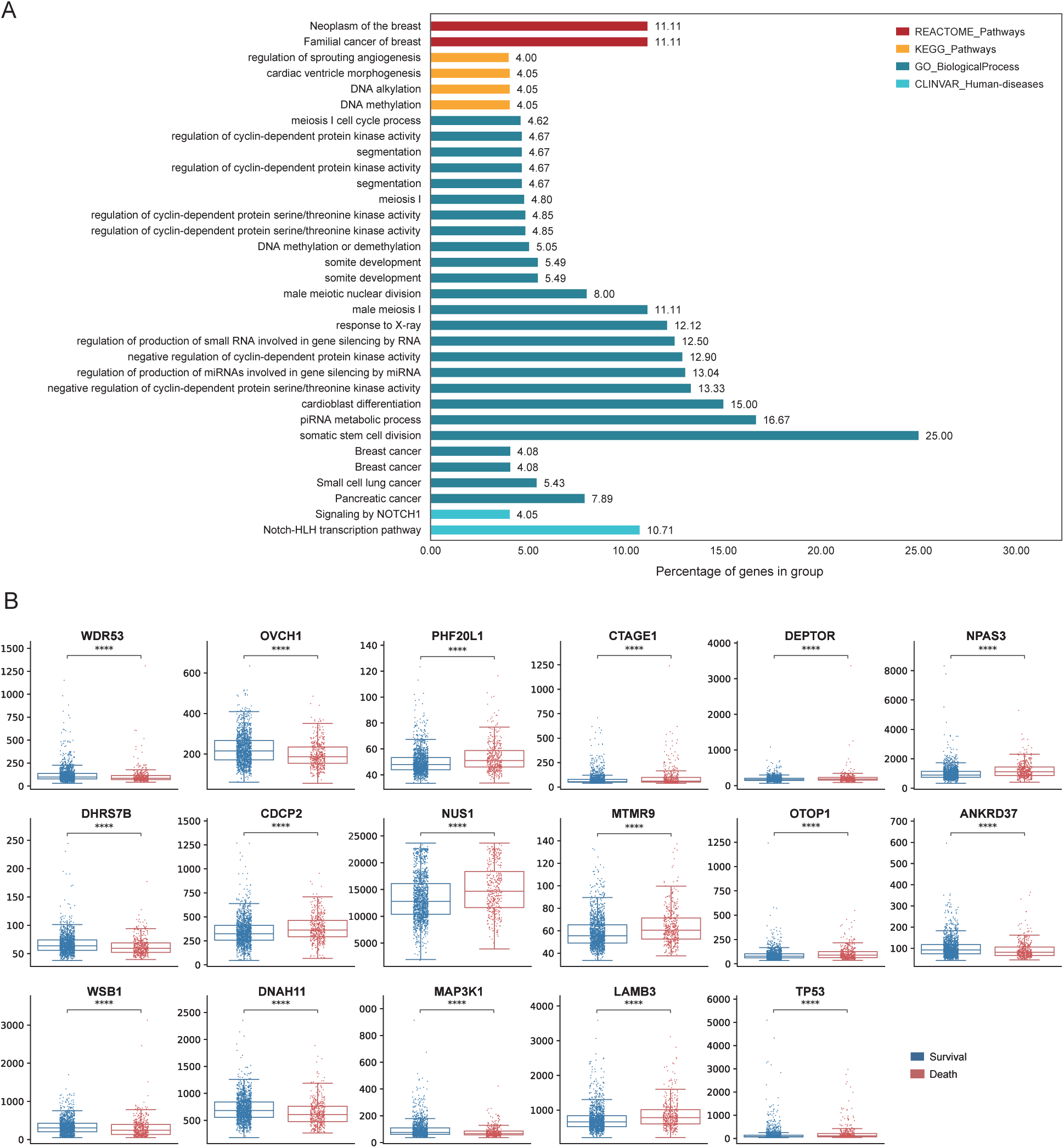
**A)** Optimal Model Pathway Enrichment Analysis; **B)** Transcriptome-level differential analysis of optimal model features.

### Difference analysis of features at the transcriptoional level

In the optimal model, we extracted the genes corresponding to 170 features (excluding two phenotypic features). In BRCA_METABRIC, a total of 17 genes were differentially expressed between the living and dead breast cancer patient groups (adjusted P value < 0.05 for all cases, Kruskal-Wallis test), such as TP53, DNAH11, MAP3K1, PHF20L1, etc. (Figure 4.B). TP53 (No. 3), DNAH11 (No. 5), MAP3K1 (No. 12), PHF20L1 (No. 20) ranked in the top 30 of the model feature weights. The limma results showed that none of these genes had a significant fold change in expression, between the living and dead breast cancer patient groups.

### Results of risk stratification

According to the model prediction results of all samples, we assigned samples with probability values less than 0.1(TPR>0.93) to the low-risk group (1473 samples), samples with probability values greater than 0.9(TNR>0.99) to high-risk group (363 samples), and others to medium-risk group (202 samples). The stratification results are shown in Figure 5.A. The Kaplan-Meier survival curves corresponding to the three sets of results are shown in Figure 5.B. The results showed that the three groups of patients had significantly different survival outcomes. This has clinical implications. For high-risk patients, more clinical intervention and active treatment may be required.

**Figure 5.**
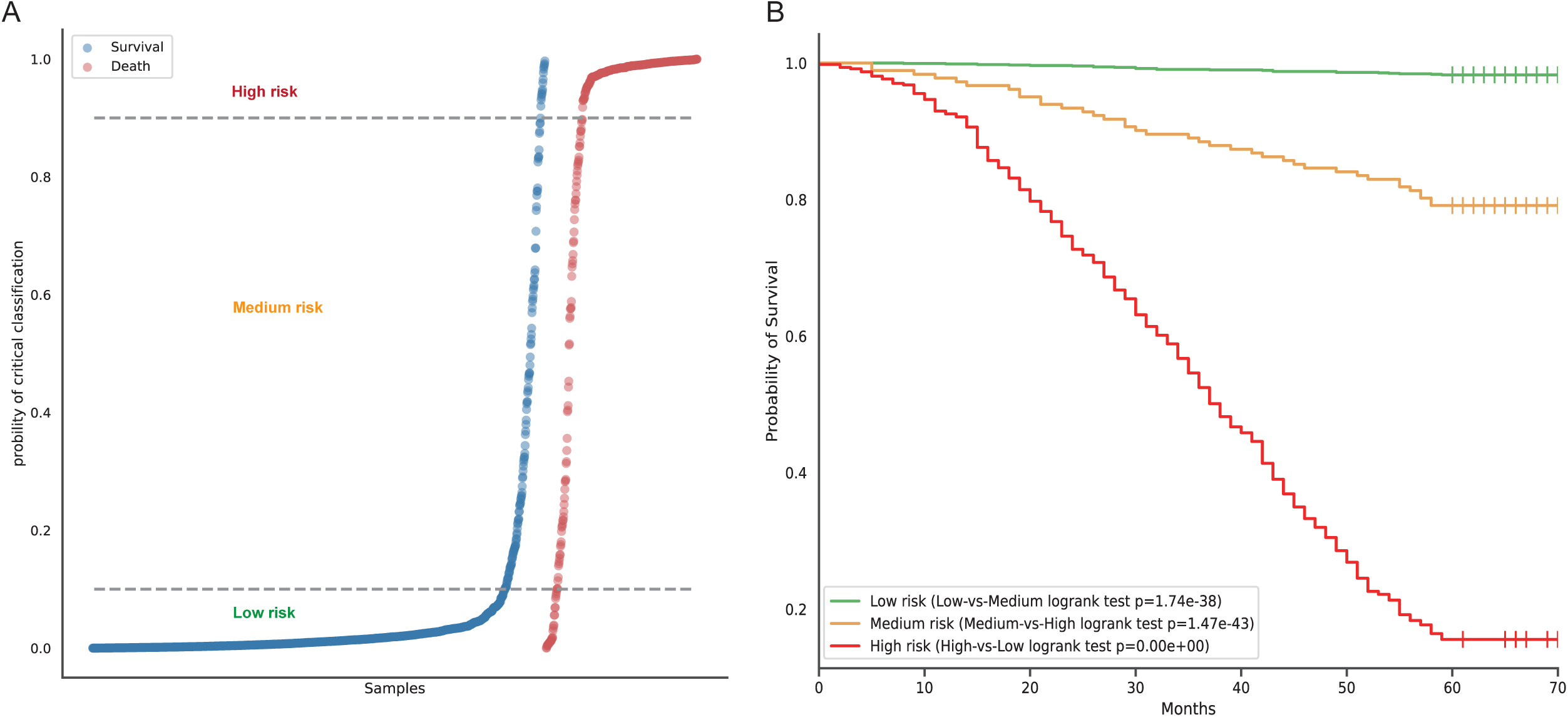
**A)** Risk stratification for all samples based on model scoring; **B)** Plot Kaplan-Meier survival curves for three groups of stratified outcomes (high, intermediate, and low risk).

## Discussion

### Model evaluation

The CatBoost algorithm model is used. Different models performed similarly in the training set and the test set without serious overfitting and strong generalization ability. The best model result is the combined variants+phenotype (AUC: 0.70). For a single feature such as SNV, CNGM has a lower AUC In an independent external data set (SNV: AUC=0.53, CNGM: AUC=0.54). CNA, as a single-dimensional feature, is similar to the combined variants model’s result in external data set(AUC=0.62), and compared to CNGM and SNV, CNA has better generalization capabilities. But this is also related to the small number of CNA and SNV features, and more comprehensive data needs to be collected for further verification. In addition, the addition of phenotype, especially age and the number of lymph nodes, has a very large impact on death, as can be seen from the feature weights of the optimal model (Figure 3).

In general, we comprehensively assessed the impact of different characteristics on the five-year mortality risk. In the process of model evaluation, we found that a single feature has poorer performance than the feature fusion model. CNA accounts for a relatively large number of model features due to the large number of original features, but there are still more CNGM features in the top 30 features. The contribution of SNV features in risk prediction is low. The addition of phenotypic information such as age and number of lymph nodes can increase the accuracy of the model.

### Discovery of biomarkers associated with five-year mortality risk

In the optimal combined variants+phenotype model, in addition to phenotypic features, some genomics features that have a greater contribution to the model have also been found. And these features still have significant differences between the survival and death groups at the transcriptome level, although there is no large fold difference. For example, TP53, DNAH11, MAP3K1, PHF20L1. As a very complex biomarker, TP53 acts as a tumor suppressor in many tumor types; induces growth arrest or apoptosis depending on the physiological circumstances and cell type which has been widely reported. Its mutations are widely present in various cancers[38-41]. IARC TP53 Database (https://p53.iarc.fr/) records all the resources of TP53 mutations[41]. They pointed out that there are 28 mutations that lead to a poor prognosis (https://p53.iarc.fr/SomaticPrognosisStats.aspx). In our model, the TP53 feature comes from the CNGM feature dimension. The model results indicate that the greater the cumulative number of TP53 mutations, the greater the probability of death within five years (Figure. 3). The DNAH11 gene mutation rs2285947 is considered a potential risk factor for ovarian cancer and breast cancer[42], and there is no clear report related to prognosis. MAP3K1 is a component of a protein kinase signal transduction cascade, which has dual regulatory effects on cell survival and apoptosis, and its regulatory mechanism is not yet clear[43, 44]. These characteristics have been reported to be related to cancer, and our study further verified their relationship with the five-year mortality risk.

Through feature selection and multi-dimensional feature fusion, the optimal model features are concentrated on pathways related to cancer, cell division, and proliferation without adding additional prior information. This reflects that the design of the model is relatively reliable, and the model can eliminate features that are not related to the training target from a large amount of input data. The genes corresponding to the features retained by the model are potential biomarkers for prognostic analysis and drug development.

## Conclusion

In general, in this article, based on the CatBoost algorithm, we use independent data sets of BRCA_METABRIC and BRCA_TCGA to conduct systematic model training on features of different dimensions. The effects of different dimensional features at the genome level on the prediction results of the model are compared. Our best model combines all the features, and the AUC in the external independent BRCA_TCGA is 0.70. In addition, the risk stratification results of all samples showed significant differences between different populations. For high-risk groups classified by the model, active clinical treatment is very necessary. This is the first five-year breast cancer death analysis based on genomic data and using external independent data for evaluation. And compared with other studies, the model based on somatic genomic variants data and phenotypic data (age, number of lymph nodes) is more prospective, and the patient’s condition can be evaluated before clinical intervention, providing guidance for follow-up treatment

Nevertheless, the research still has limitations. When selecting the features that the two data sets contain in common, the SNP and CNGM features only get very little intersection, which may lead to the underestimation of the role of SNP and CNGM. Deep learning algorithms have not been used and compared. We will continue to conduct in-depth research, collect more comprehensive data, design and develop new algorithms based on existing experience, and further compare the performance differences between machine learning and deep learning. In addition, we will also try to collect other cancer data, conduct migration learning, and develop a five-year mortality risk model for pan-cancer.

## Acknowledgments

Thanks to all researchers and subjects who volunteered to participate in this project.

## Supplementary Figure

**Supplementary Figure S1**. Statistical results of sample distribution regarding gender, number of lymph nodes, menopause (−9 - unknown, 0 - not menopause, 1 - menopause).

**Supplementary Figure S2**. The optimization process of the five models (CNA, SNV, CNGM, SNV+CNGM+CNA(combined variants), combined variants+phenotype).

## Reference

1. Sung, H., et al., Global cancer statistics 2020: GLOBOCAN estimates of incidence and mortality worldwide for 36 cancers in 185 countries. CA: a cancer journal for clinicians, 2021. 71(3): p. 209–249.

2. Loibl, S., et al., Breast cancer. Lancet, 2021.

3. Garcia-Closas, M., et al., Heterogeneity of breast cancer associations with five susceptibility loci by clinical and pathological characteristics. PLoS genetics, 2008. 4(4): p. e1000054.

4. Lüönd, F., S. Tiede, and G. Christofori, Breast cancer as an example of tumour heterogeneity and tumour cell plasticity during malignant progression. British Journal of Cancer, 2021. 125(2): p. 164–175.

5. McClellan, J. and M.-C. King, Genetic heterogeneity in human disease. Cell, 2010. 141(2): p. 210–217.

6. Allemani, C., et al., Global surveillance of trends in cancer survival 2000-14 (CONCORD-3): analysis of individual records for 37 513 025 patients diagnosed with one of 18 cancers from 322 population-based registries in 71 countries. Lancet, 2018. 391(10125): p. 1023–1075.

7. Haybittle, J.L., et al., A prognostic index in primary breast cancer. British Journal of Cancer, 1982. 45(3): p. 361–366.

8. Cox, D.R., Regression Models and Life-Tables. Journal of the Royal Statistical Society. Series B (Methodological), 1972. 34(2): p. 187–220.

9. Kattan, M.W., et al., A tool for predicting breast carcinoma mortality in women who do not receive adjuvant therapy. Cancer, 2004. 101(11): p. 2509–15.

10. Ravdin, P.M., et al., Computer program to assist in making decisions about adjuvant therapy for women with early breast cancer. J Clin Oncol, 2001. 19(4): p. 980–91.

11. Campbell, H.E., et al., Estimation and external validation of a new prognostic model for predicting recurrence-free survival for early breast cancer patients in the UK. Br J Cancer, 2010. 103(6): p. 776–86.

12. Engelhardt, E.G., et al., Accuracy of the online prognostication tools PREDICT and Adjuvant! for early-stage breast cancer patients younger than 50 years. Eur J Cancer, 2017. 78: p. 37–44.

13. Reis-Filho, J.S. and L. Pusztai, Gene expression profiling in breast cancer: classification, prognostication, and prediction. Lancet, 2011. 378(9805): p. 1812–23.

14. van ‘t Veer, L.J., et al., Gene expression profiling predicts clinical outcome of breast cancer. Nature, 2002. 415(6871): p. 530–6.

15. Paik, S., et al., A multigene assay to predict recurrence of tamoxifen-treated, node-negative breast cancer. N Engl J Med, 2004. 351(27): p. 2817–26.

16. Filipits, M., et al., A new molecular predictor of distant recurrence in ER-positive, HER2-negative breast cancer adds independent information to conventional clinical risk factors. Clin Cancer Res, 2011. 17(18): p. 6012–20.

17. Manjang, K., et al., Prognostic gene expression signatures of breast cancer are lacking a sensible biological meaning. Sci Rep, 2021. 11(1): p. 156.

18. Kourou, K., et al., Machine learning applications in cancer prognosis and prediction. Comput Struct Biotechnol J, 2015. 13: p. 8–17.

19. Hieronymus, H., et al., Tumor copy number alteration burden is a pan-cancer prognostic factor associated with recurrence and death. Elife, 2018. 7.

20. Nguyen, C., Y. Wang, and H.N. Nguyen, Random forest classifier combined with feature selection for breast cancer diagnosis and prognostic. Journal of Biomedical Science and Engineering, 2013. Vol.06 No.05: p. 10.

21. Park, K., et al., Robust predictive model for evaluating breast cancer survivability. Engineering Applications of Artificial Intelligence, 2013. 26(9): p. 2194–2205.

22. Zhang, Y., et al., Toward the precision breast cancer survival prediction utilizing combined whole genome-wide expression and somatic mutation analysis. BMC Med Genomics, 2018. 11(Suppl 5): p. 104.

23. He, Z., et al., Integrating Somatic Mutations for Breast Cancer Survival Prediction Using Machine Learning Methods. Front Genet, 2020. 11: p. 632901.

24. Li, J., et al., Predicting breast cancer 5-year survival using machine learning: A systematic review. PLoS One, 2021. 16(4): p. e0250370.

25. Sun, D., M. Wang, and A. Li, A multimodal deep neural network for human breast cancer prognosis prediction by integrating multi-dimensional data. IEEE/ACM Trans Comput Biol Bioinform, 2018.

26. Arya, N. and S. Saha, Multi-modal advanced deep learning architectures for breast cancer survival prediction. Knowledge-Based Systems, 2021. 221: p. 106965.

27. Tomczak, K., P. Czerwińska, and M. Wiznerowicz, The Cancer Genome Atlas (TCGA): an immeasurable source of knowledge. Contemporary oncology, 2015. 19(1A): p. A68.

28. Curtis, C., et al., The genomic and transcriptomic architecture of 2,000 breast tumours reveals novel subgroups. Nature, 2012. 486(7403): p. 346–352.

29. Pereira, B., et al., The somatic mutation profiles of 2,433 breast cancers refine their genomic and transcriptomic landscapes. Nature communications, 2016. 7(1): p. 1–16.

30. Rueda, O.M., et al., Dynamics of breast-cancer relapse reveal late-recurring ER-positive genomic subgroups. Nature, 2019. 567(7748): p. 399–404.

31. Kraskov, A., H. Stögbauer, and P. Grassberger, Estimating mutual information. Physical review E, 2004. 69(6): p. 066138.

32. Li, F. and Y. Yang. Analysis of recursive feature elimination methods. in Proceedings of the 28th annual international ACM SIGIR conference on Research and development in information retrieval. 2005.

33. Kursa, M.B. and W.R. Rudnicki, Feature selection with the Boruta package. J Stat Softw, 2010. 36(11): p. 1–13.

34. Lobo, J.M., A. Jimenez-Valverde, and R. Real, AUC: a misleading measure of the performance of predictive distribution models. Global Ecology and Biogeography, 2008. 17(2): p. 145–151.

35. Lundberg, S.M. and S.-I. Lee. A unified approach to interpreting model predictions. in Proceedings of the 31st international conference on neural information processing systems. 2017.

36. Bindea, G., et al., ClueGO: a Cytoscape plug-in to decipher functionally grouped gene ontology and pathway annotation networks. Bioinformatics, 2009. 25(8): p. 1091–3.

37. Smyth, G.K., Limma: linear models for microarray data, in Bioinformatics and computational biology solutions using R and Bioconductor. 2005, Springer. p. 397–420.

38. Bertheau, P., et al., TP53 status and response to chemotherapy in breast cancer. Pathobiology, 2008. 75(2): p. 132–139.

39. BørresenLJDale, A.L., TP53 and breast cancer. Human mutation, 2003. 21(3): p. 292–300.

40. Petitjean, A., et al., TP53 mutations in human cancers: functional selection and impact on cancer prognosis and outcomes. Oncogene, 2007. 26(15): p. 2157–2165.

41. Petitjean, A., et al., Impact of mutant p53 functional properties on TP53 mutation patterns and tumor phenotype: lessons from recent developments in the IARC TP53 database. Human mutation, 2007. 28(6): p. 622–629.

42. Verma, S., et al., Genetic variants of DNAH 11 and LRFN 2 genes and their association with ovarian and breast cancer. International Journal of Gynecology & Obstetrics, 2020. 148(1): p. 118–122.

43. Pham, T.T., S.P. Angus, and G.L. Johnson, MAP3K1: genomic alterations in cancer and function in promoting cell survival or apoptosis. Genes & cancer, 2013. 4(11-12): p. 419–426.

44. Xue, Z., et al., MAP3K1 and MAP2K4 mutations are associated with sensitivity to MEK inhibitors in multiple cancer models. Cell research, 2018. 28(7): p. 719–729.

