## Supplementary Figure for "Machine learning enables accurate prediction of breast cancer five-year survival using somatic genomic variants"

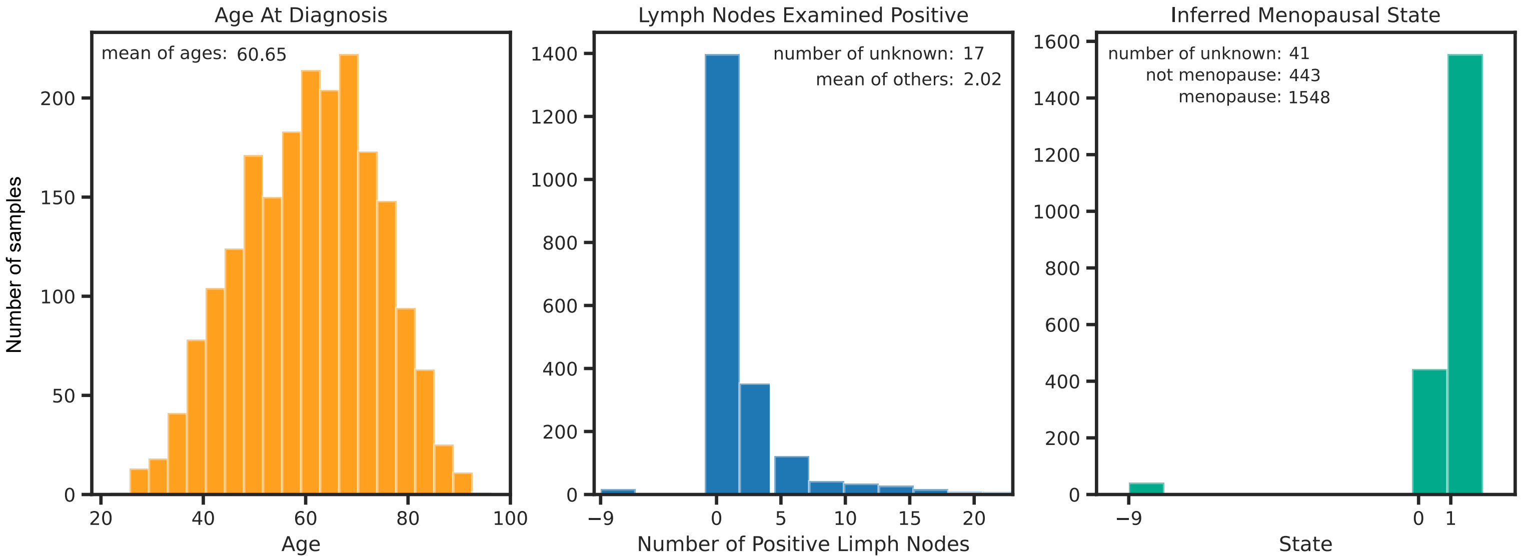


**Supplementary Figure S1.** Statistical results of sample distribution regarding gender, number of lymph nodes, menopause (-9 - unknown, 0 - not menopause, 1 - menopause).

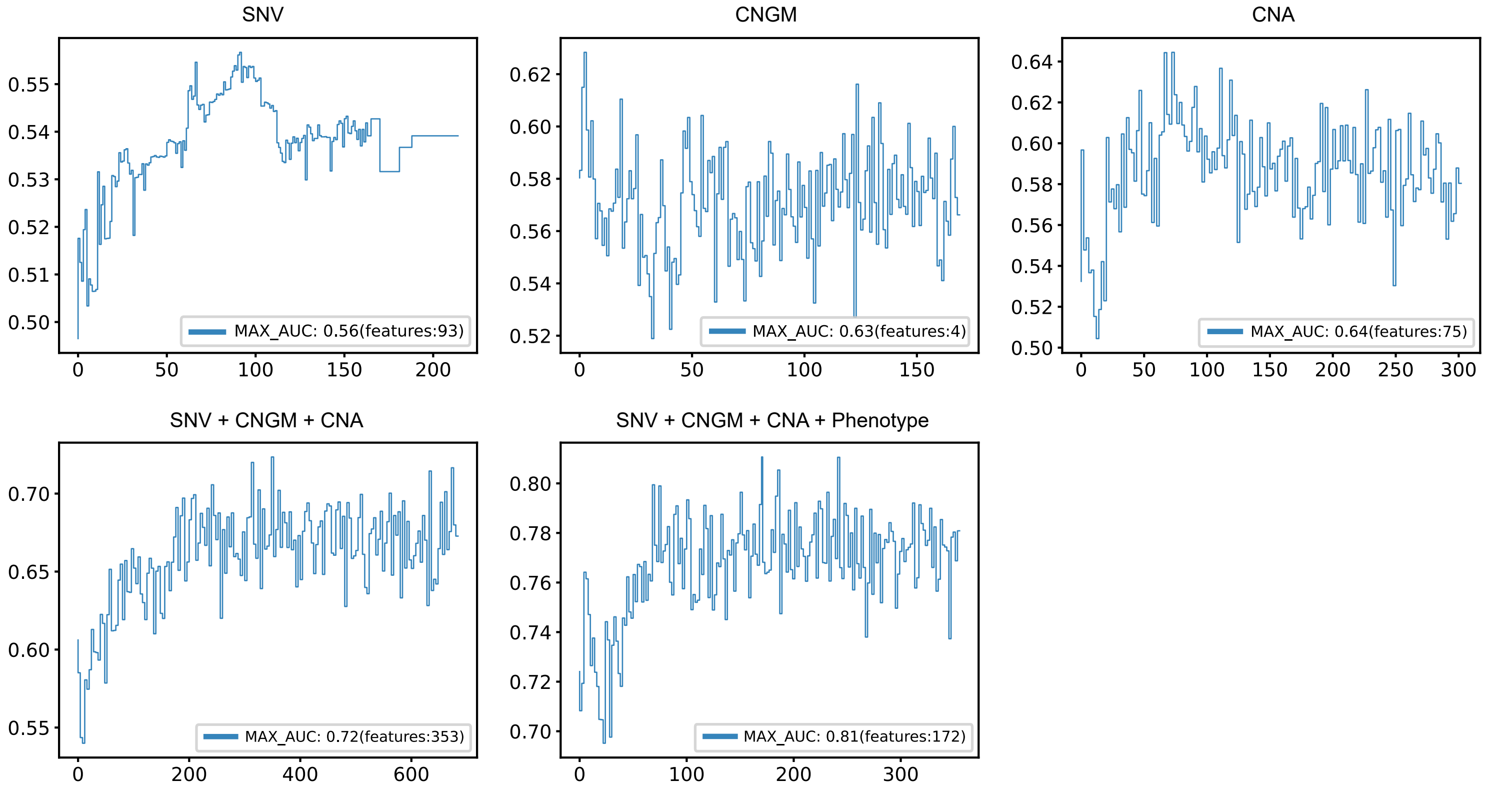


**Supplementary Figure S2.** The optimization process of the five models (CNA, SNV, CNGM, SNV+CNGM+CNA(combined variants) , combined variants+phenotype).
